# Molecular Dynamics Simulation of the *E.coli* FtsZ

**DOI:** 10.1101/280222

**Authors:** Vidyalakshmi C Muthukumar

## Abstract

Previous molecular dynamics studies of the FtsZ protein revealed that the protein has high intrinsic flexibility which the crystal structures were unable to reveal. The initial configuration in these studies was based on the available crystal structure data and therefore, the effect of the C-terminal Intrinsically Disordered Region (IDR) of FtsZ could not be observed in these previous studies. Recent investigations have revealed that the C-terminal IDR is crucial for FtsZ assembly *in vitro* and Z ring formation *in vivo*. Therefore, in this study, we simulate FtsZ with the IDR.

Simulations of the FtsZ monomer in different nucleotide bound forms (without nucleotide, GTP, GDP) were performed. In the conformations of FtsZ monomer with GTP, GTP binds variably with the protein. Such variable interaction with the monomer has not been observed in any previous simulation studies of FtsZ and not observed in crystal structures. The central helix bends towards the C-terminal domain in the GTP bound form, thus making way for polymerization. Nucleotide dependent small shift/rotation of the C-terminal domain was observed in average structures.

## 1. Introduction

Many crystal structures of FtsZ are available in the PDB (the Protein Data Bank). For e.g. *P. aeruginosa* and *M. jannaschii* FtsZ crystal structures (PDB ID: 2VAW [1] and 1W5A/1W5B [2]). The protein has a conserved globular core which is approximately 300 residues. This core comprises of the ordered N- and C-terminal domain residues. GTP binds to the nucleotide binding site present in N-terminal domain. The protein also contains a short N-terminal intrinsically disordered peptide (IDP) and a much longer C-terminal intrinsically disordered region (IDR). The N-terminal IDP is present in the crystal structure of the *P. aeruginosa* FtsZ 2VAW. The IDR has not been resolved by X-ray crystallization. This region is not conserved and its length may vary considerably across different bacterial species [3]. However, the extreme C-terminus consists of a conserved ∼ 11 residue C-terminal helix which interacts with regulatory proteins like ZipA. The crystal structure of the extreme C-terminal residues 375 – 383 of *E. coli* FtsZ with ZipA is available (PDB ID: 1F47 [4]).

In a previous study, FtsZ crystals obtained from different species were compared [1]. Root mean square deviation (RMSD) between the available structures and rotation angle of the C-terminal domain were calculated. The RMSD or difference in rotation angles were not found to be significant for the possibility of different FtsZ conformations to be present in these crystal structures. The conclusion which was drawn from the comparison was that there could be a possibility of inter-species differences in the crystal structures, however, the existence of nucleotide dependent different conformations of FtsZ was unlikely.

Using electron microscopy, FtsZ protofilaments have been visualized. It is observed that FtsZ protofilaments may be straight or bent depending on nucleotide binding. *In vitro* polymerization studies support a more straight conformation of the GTP bound protofilament and curved GDP bound protofilaments [5].

Molecular dynamics simulations support similar ideas. Significant conformational changes in the monomer as such, were not observed in simulations [6, 7]. However, the studies definitely revealed the high intrinsic flexibility of the protein. In FtsZ monomer simulations Kathiresan *et al.* [6], reported a ‘twisted bending’ of the core helix and the possibility that the ‘H6 – H7’ region (top portion of the central helix) acts a switch which recognizes the nucleotide binding state of the protein. Principal component analysis (PCA) revealed that in the GTP simulations, the top portion of the central helix moves upwards (towards the C-terminal domain of the top subunit in the longitudinal protofilament) and the HC2, HC3 of C-terminal domain moves downwards to make protofilament longitudinal contacts. Even though simulations reveal important nucleotide dependent structural changes in the monomer, conformational change in the monomer is not observed. RMSD between the GTP and GDP forms are very low, comparable to crystal structure resolution. In FtsZ dimer simulations [7], the nucleotide dependent change in protofilament curvature was attributed to the loss of monomer-monomer contact only.

Recent studies have brought to light the role of the C-terminal linker (CTL) (C-terminal IDR except the extreme C-terminal conserved helix) in Z-ring assembly [8, 9]. Removal of the CTL, in *B. subtilis* leads to a filamentous phenotype in which Z-ring assembly does not occur. *In vitro,* the removal of CTL resulted in a much lower GTPase activity (2.5 fold lower GTPase) and an increase in FtsZ critical concentration (by 1.5 fold) and inhibited protofilament formation. It was observed that intrinsic flexibility is crucial for FtsZ function, and replacement with rigid alpha-helix hinders FtsZ assembly *in vitro* and cell division *in vivo*. It was observed that ‘FtsZ CTLH’ (FtsZ in which the CTL had been replaced by rigid helical repeats) could form only small oligomers *in vitro.* CTLs of varying lengths (25 -100 residues) were functional *in vivo.* Since CTL is required for FtsZ assembly *in vitro* (in the absence of regulatory proteins), therefore, the CTL might be playing an important role during the formation of protofilaments.

In the present study, we study the role of the FtsZ IDR using molecular dynamics. Simulations were performed using a homology model for the *E. coli FtsZ* in which the C-terminal IDR was included. The wild-type monomer was simulated without nucleotide, with GTP or with GDP (guanosine diphosphate). In all the simulations, it was observed that the globular core maintains its structure and the IDR is very flexible. Differential flexibilities of the IDR were observed in the wild-type monomer simulations with GDP and GTP. We also significant bending of the central helix in GTP simulations and the possibility of C-terminal domain rotation.

## 2. Methods

### 2.1 Homology Model of the *E. coli* FtsZ protein (wild-type, WT)

For constructing a model of the WT *E. coli* FtsZ protein, the gene: ftsZ (EC2860050_0094) was identified (NCBI accession is <u>AQDL01000001.1</u>, locus tag is EC2860050_0094). The primary sequence was obtained using the ExPASy Translate tool [10]. The obtained primary sequence was aligned with primary sequences of FtsZ proteins from other bacterial species, for which crystal structures are already available in the Protein Data Bank. BLAST-p [11] alignment scores are provided in Table 2.1.

**Table 2.1.** Alignment of E. coli FtsZ protein (obtained by translating Gene: ftsZ (EC2860050_0094)) with various subject sequences using the Standard Protein Blast algorithm.

| Species and Transcript ID | Max. Score | Total Score | Query Cover (%) | E-value | Ident. (%) | Positives (%) | No. of Matches |
| --- | --- | --- | --- | --- | --- | --- | --- |
| *P. aeruginosa*<br>PAM18_4498 | 392 | 392 | 93 | 4e-138 | 57 | 69 | 1 |
| *M. tuberculosis*<br>MTCTRI2_2186 | 253 | 253 | 93 | 3e-84 | 45 | 63 | 1 |
| *B. subtilis*<br>BEST7613_2297 | 279 | 298 | 98 | 8e-94 | 47 | 64 | 2 |
| *T. maritima* Tmari_0838 | 254 | 269 | 82 | 6e-85 | 48 | 66 | 2 |
| *A. aeolicus* aq_525 | 231 | 231 | 77 | 6e-76 | 46 | 63 | 1 |
| *S. aureus*<br>SA21334_0269 | 270 | 285 | 81 | 9e-91 | 51 | 72 | 2 |
| *S. epidermidis*<br>SEVCU045_1562 | 266 | 266 | 91 | 3e-89 | 45 | 64 | 1 |
| *H. volcanii* ADE02725 | 230 | 244 | 81 | 2e-75 | 45 | 65 | 2 |

The *P. aeruginosa* FtsZ gave the largest total score, query cover and highest E-value. Therefore, the crystal structure 2VAW of *P. aeruginosa* FtsZ was selected as the primary template for homology modelling. *E.coli* FtsZ protein sequence is 383 amino acids. In all available FtsZ crystal structures including 2VAW, structural information is available for the ordered residues of the globular core only. In 2VAW, in addition, the short disordered peptide at the N-terminal is also present. Therefore, this structure was used as a template for residues 2 – 316 of the *E. coli* FtsZ protein *i.e.* the residues which align in the two sequences and for which structural information is available in the 2VAW FtsZ molecule.

However, the IDR at the C-terminal is not resolved in any of the crystal structures. Since, residues 2 – 316 of *E. coli* FtsZ align with residues 2 – 316 of the *P. aeruginosa* FtsZ sequence, therefore, it may be assumed that residues 317 – 383 in *E. coli* FtsZ correspond to the C-terminal IDR (including the extreme C-terminal conserved helix).

On analysing the primary sequence (PrDOS probability described later) it was observed that the residues of the IDR have the highest disorder probability. Therefore, during molecular dynamics simulation, significant conformational changes/rearrangements in this region were expected. We selected templates for this region by aligning short sequences from the IDR with other proteins in the PDB database. PDB Extended Blast-P search (E. value cut off up to 10) for *E. coli* FtsZ query sequence gave an MP1-p14 complex (PDB accession: 1SKO, molecule: Mitogen-activated protein kinase kinase 1 interacting protein 1, chain A: 143 amino acids, [12]). A 41 residue sequence in this protein, from 21 – 61 (crystal residues), aligned with residues 342 – 382 in *E. coli* FtsZ. 48% positive alignment was observed for this sequence. For building homology model, residues 21 – 62 of the 1SKO structure were used as template for residues 342 – 383 of *E. coli* FtsZ. To identify suitable templates for residues 317 – 341, shorter peptides from the C-terminal flexible / disordered region were used as query sequences. For the query sequence 325 – 360 in *E. coli* FtsZ, one of the results was residues 172 – 207 in *Marinibacillus marinus* Adenylate kinase protein, PDB accession 3FB4 [13]. For residues, 317–324 also, we used neighbouring residues from the 3FB4 sequence. Therefore, including additional 11 neighbouring residues, residues 161 – 207 of the 3FB4 structure was used as template for residues 317 – 360 of *E. coli* FtsZ (48% positive alignment, included 3 gaps in the match). The C-terminal residues 367 – 383 of the *E. coli* FtsZ protein has already been crystallized with the protein ZipA [4]. It was used as a template for the corresponding residues. The templates used for constructing *E. coli* FtsZ homology model are listed in Table 2.2. Modeller [14 – 16] program was used for building the homology model. One model was generated. The model generated includes residues Phenylalanine (residue no. 2) – Aspartate (residue no. 383), Figure 2.1.

**Table 2.2.** Template crystal structures for constructing *E. coli* FtsZ homology model.

| S. No. | PDB | Protein | Aligned residues from *E. coli* FtsZ |
| --- | --- | --- | --- |
| 1 | 2VAW [1] (2 – 316) | *P. aeruginosa* FtsZ | 2 – 316 |
| 2 | 1SKO [12] (A:21 – 62) | *Homo sapiens* Mp1-p14 kinase | 342 – 383 |
| 3 | 3FB4 [13] (A:161 – 207) | *M. marinus* Adenylate kinase | 317 – 360 |
| 4 | 1F47 [4] (A:1 – 17) | ZipA-FtsZ *E. coli* | 367 – 383 |

**Fig. 2.1).**
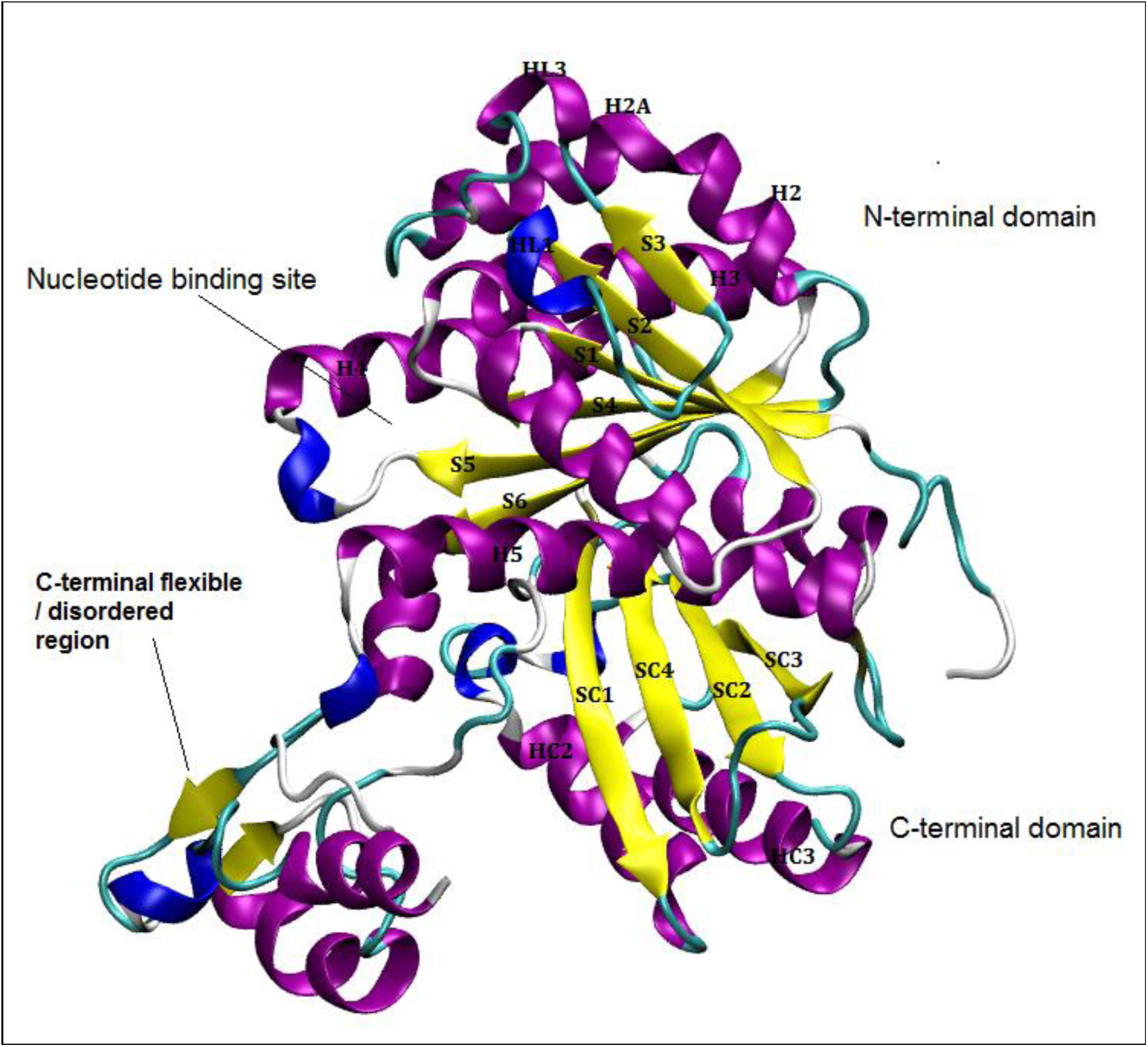
*E. coli* FtsZ model built using templates given in Table 2.3.

Coordinates for nucleotides GDP, GTP were obtained from the Hetero-compound Information Centre, Uppsala. Nucleotides were allowed to bind freely by placing inside the nucleotide binding site or close to it. Initial position of the nucleotides is provided in Fig. 2.2. Two independent repeat simulations were performed with GDP. Five independent repeat simulations were performed with GTP using two different starting orientations of the nucleotide.

**Fig. 2.2).**
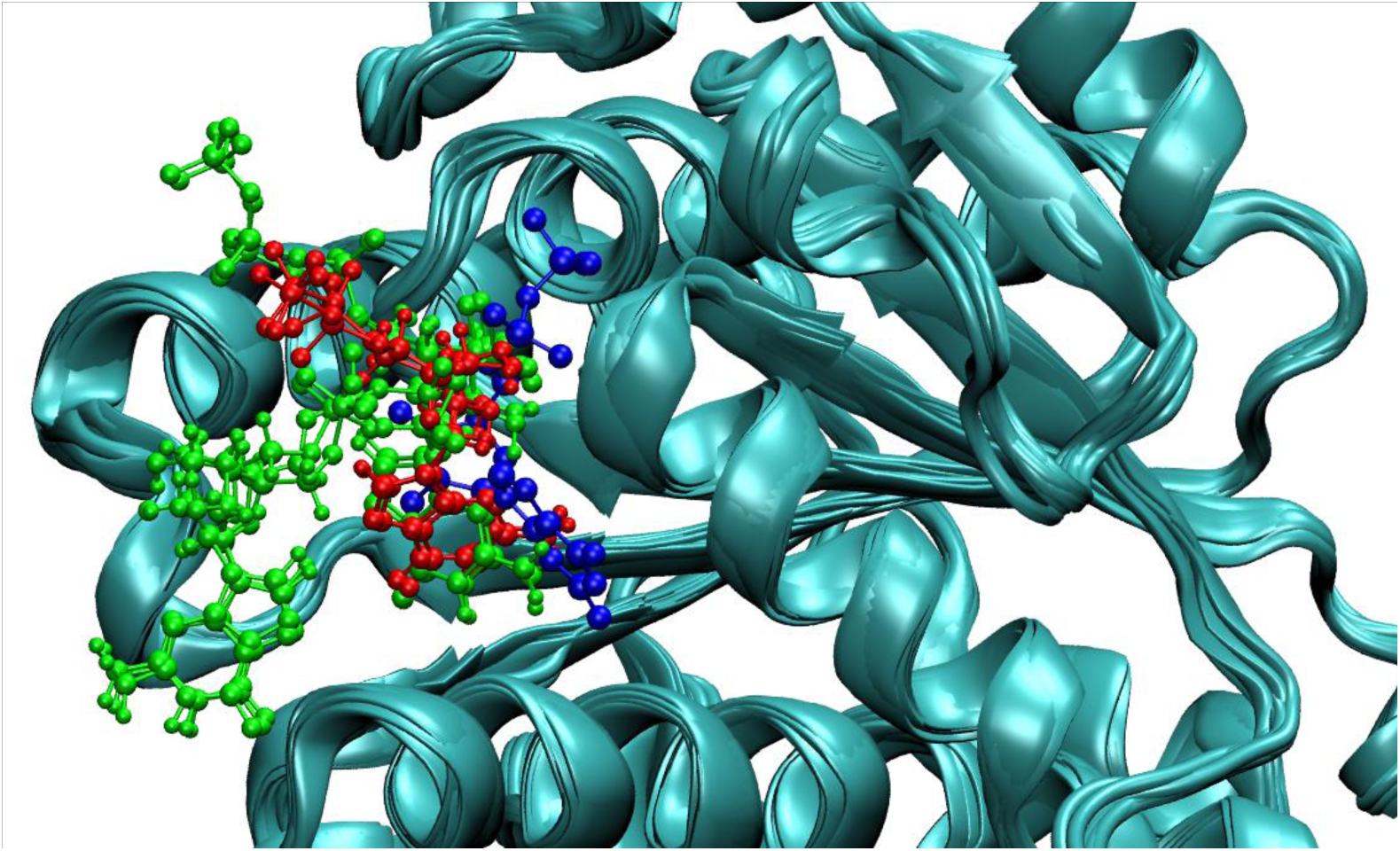
Position of nucleotides in NPT equilibrated structures GDP in 2VAW reference (blue), GDP in 2 simulations (red), 2 different orientations of GTP in 5 simulations (green).

### 2.2 Molecular dynamics simulations

Atomistic molecular dynamics simulations were performed using GROMACS version 4.6.5 [17]. Simulations were performed in the AMBER94 force field [18, 19]. AMBER parameters for nucleotides developed by Meagher et al. [20] was used. Protein molecule was centred in the cubic box, at a minimum distance of 1.5 nm from box edges. Solvent (water) molecules were added in the coordinate file. SPC-E (Single Point Charge Extended) water model configuration was used [21]. Mg^2+^ and Cl^−^ ions were added to neutralize the simulation box and at a minimum concentration of up to 10 mM (any of which was higher). Energy minimization was performed using the steepest descent minimization algorithm until the maximum force on any atom in the system was less than 1000.0 kJ/mol/nm. A step size of 0.01 nm was used during energy minimization. A cut-off distance of 1 nm was used for generating the neighbour list and this list was updated at every step. Electrostatic forces were calculated using Particle-Mesh Ewald method (PME) [22]. A cut-off of 1.0 nm was used for calculating electrostatic and Van der Waals forces. Periodic boundary conditions were used. A short 100 ps NVT equilibration was performed. During equilibration, the protein molecule was restrained. Leap-frog integrator MD simulation algorithm [23] was implemented with a time-step of 2 fs. All bonds were constrained by the LINear Constraints Solver (LINCS) constraint-algorithm [24]. Neighbour list was updated every 10 fs. A distance cut-off of 1 nm was used for calculating both electrostatic and van der Waals forces. Electrostatic forces were calculated using PME method. Two groups *i.e.* protein and non-protein (solvent, ligand and ions) were coupled with the modified Berendsen thermostat [25] set at 300 K. The time constant for temperature coupling was set at 0.1 ps. Long range dispersive corrections were applied for both energy and pressure. Coordinates were saved either every 2 ps or every 5 ps (in .xtc compressed trajectory format). A short 100 ps NPT equilibration was performed similar to the NVT equilibration with Parrinello-Rahman barostat [26, 27], with time constant of 2 ps, was applied to maintain pressure at a constant value of 1 bar. 100 ns MD simulation in NPT ensemble was implemented.

## 3. Results and Discussion

### 3.1 Pr**otein** D**isOrder prediction** S**ystem (PrDOS) prediction of disordered residues in the *E. coli* FtsZ protein sequence.**

Algorithms such as PrDos predict disordered regions in proteins [28]. PrDos predictions are based on (a) local amino acid sequence information and (b) alignments with homologues for which structures have been determined already. Combination (based on weighted average) of these two independent predictions gives the probability of disorder. The threshold probability above which a residue is said to be disordered is 0.5. For the *E. coli* FtsZ protein sequence, the stretch of residues 319 – 368 was predicted as a disordered region in the protein (2 state prediction results in Fig. 3.1). The short N-terminal peptide of ∼ 8 to 10 residues, is found to be in a helical conformation in some FtsZ molecules (e.g. 1W5A), in a loop (e.g. 2VAW) or could not be resolved in some structures (e.g. 2R6R). According to PrDOS predictions these residues, residues 1 – 8, also have disorder probabilities greater than 0.5.

**Fig. 3.1).**
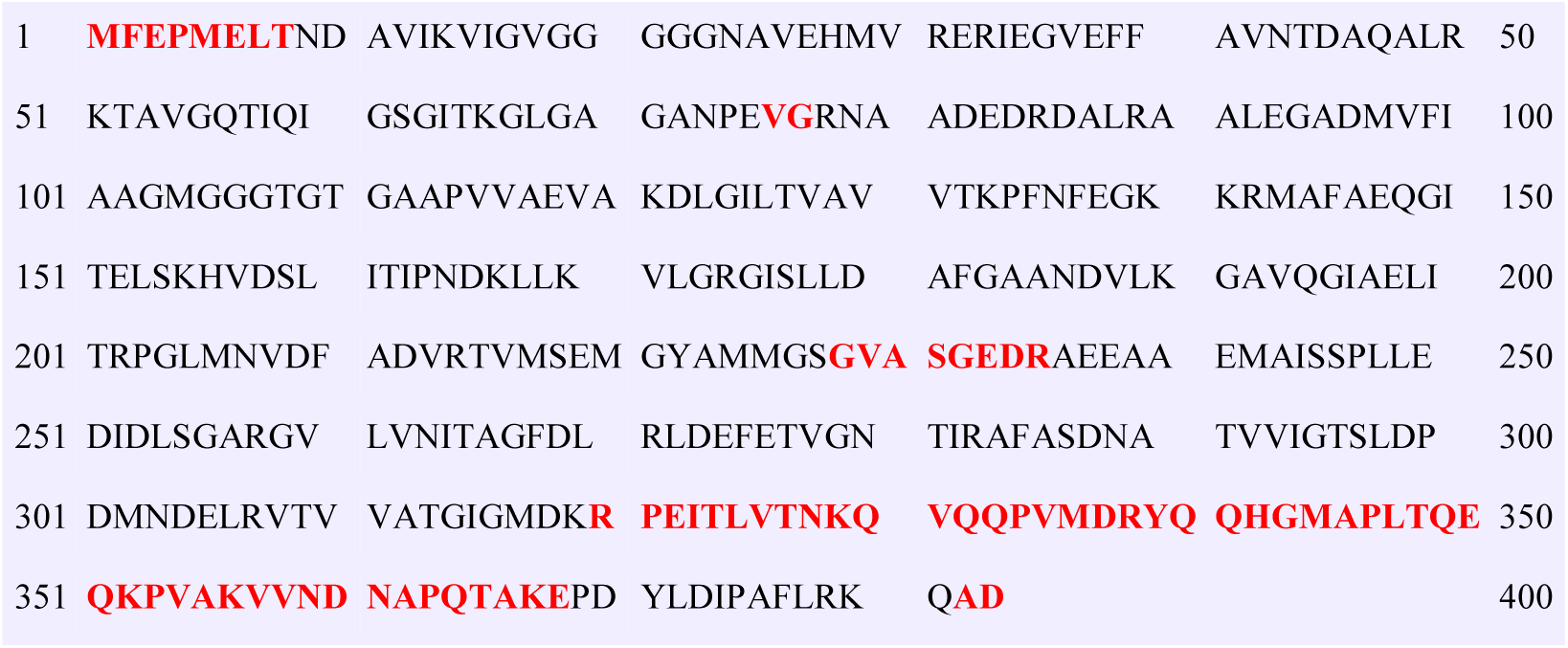
PrDos 2 state prediction (False positive rate 5%) Red: Disordered residues, Black: Ordered residues.

### 3.2 Assessment of the model

The RMSD of the model and the 2VAW template based on backbone alignment was 3.3 Å. The model for the IDR was built using the templates 3FB4 residues 161 –207 for residues 317–360 of *E.coli* FtsZ. The Modeller predicted the residues 317 – 330 in either coil or turn conformations, 331 – 341 in an α-helix, 342 – 357 in a β-haripin loop and 358 – 362 are in either coil or turn conformations, 363 – 366 are in a 3-10 helix, 367 – 371 are in a turn conformation, 372 – 381 are in an α-helix and 382 in a coil. The model was assessed using ProQ which generates LG and Maxsub scores based on atom-atom contacts, residue-residue contacts and solvent accessibility. LG score for the model was 5.396 (in the ProQ very good model range).

### 3.3 MD visualization

Molecular dynamics simulations were viewed in VMD. To show the evolution of the protein in simulation, structure of the protein at every 20 ns in the wild-type monomer simulation WT-FtsZ mono-GDP (1) is provided in Fig. 3.2. Structure of the protein at a single time frame from well equilibrated simulation provides considerable information about the simulation. Conformational flexibility (present in loop regions) which are lost during averaging of structures may be observed from structures at a given time in the simulation. In some of the simulations we observed a large change in curvature of the central helix of FtsZ. The extent of bending was found to differ between the independent repeat simulations of the same system. An average structure does not show the extent to which such variations occur. Therefore, for the discussion, we have used representative structures obtained from well equilibrated simulations. We use structures from the same time frame 93.9 ns from 100 ns MD simulation (Fig. 3.3).

It may be observed that the structure of the globular core is stable. However, the structure of the C-terminal IDR is not stable and its structure varies during simulations (Fig. 3.2 a – f), and between different simulations also (Fig. 3.3 a – i), consistent with its intrinsic disorder.

In the two independent repeat simulations with GDP, WT-FtsZ mono-GDP (1) and (2), GDP is located well inside the nucleotide binding site (Fig. 3.3 c – d). Frames at 20 ns interval for the simulation WT-FtsZ mono-GDP (1) given in Fig. 3.2 (a – f). It may be observed that after t = 20 ns, there is no further significant change in the location of GDP apart from a small change in the phosphate tail between t= 80 ns (Fig. 3.2 e) to t = 100 ns (Fig. 3.2 f). High variation in position and orientation of the GTP nucleotide is observed. For e.g., in Fig. 3.3 f) GTP guanine group points towards the nucleotide binding site and its phosphate tail points outwards but in Fig. 3.3 g) it is oriented in the opposite direction. The position of GTP was observed to fluctuate continuously during the simulations suggesting that the GTP binding is unstable, probably held by non-specific interactions with the more flexible loop residues of the nucleotide binding site.

The secondary structure of residues in the region HL1 – S3 – (loop between S3 and H2A) (residue nos. 48 – 72) and S5 – H4 loop varied considerably. These residues/regions are labelled in Fig. 3.3 (a), the secondary structure elements are labelled in Fig. 3.3 (b). The residues of HL1, form a 3-10 helix (Fig. 3.3 (b), (c)), α-helix (Fig. 3.3 (d)). It may be observed that in the simulation WT-FtsZ mono-GTP (3) (Figure 3.3(g) and (i)), the short β-strand S3 converted from β-strand to a β-bridge (formed by a single residue). Residues between S3 and H2A form α-helix (Fig. 3.3 (e)), 3-10 helix (Fig. 3.3 (c)) in some simulations or present in a loop in others (Fig. 3.3 (g) and (h)). Similar secondary structure changes may be seen in the S5 H4 loop which forms a 3-10 helix in some simulations (Fig. 3.3 (c), (d), (e), (g), (h), (i)). Varying conformations of these residues of the nucleotide binding site may be essential for allowing nucleotide binding and release. Different conformations observed in repeat simulations of the same nucleotide binding state, and at different time points in the same simulation, suggests that these residues adapt to nucleotide binding and identical conformation of these residues is not necessary.

**Fig. 3.2.**
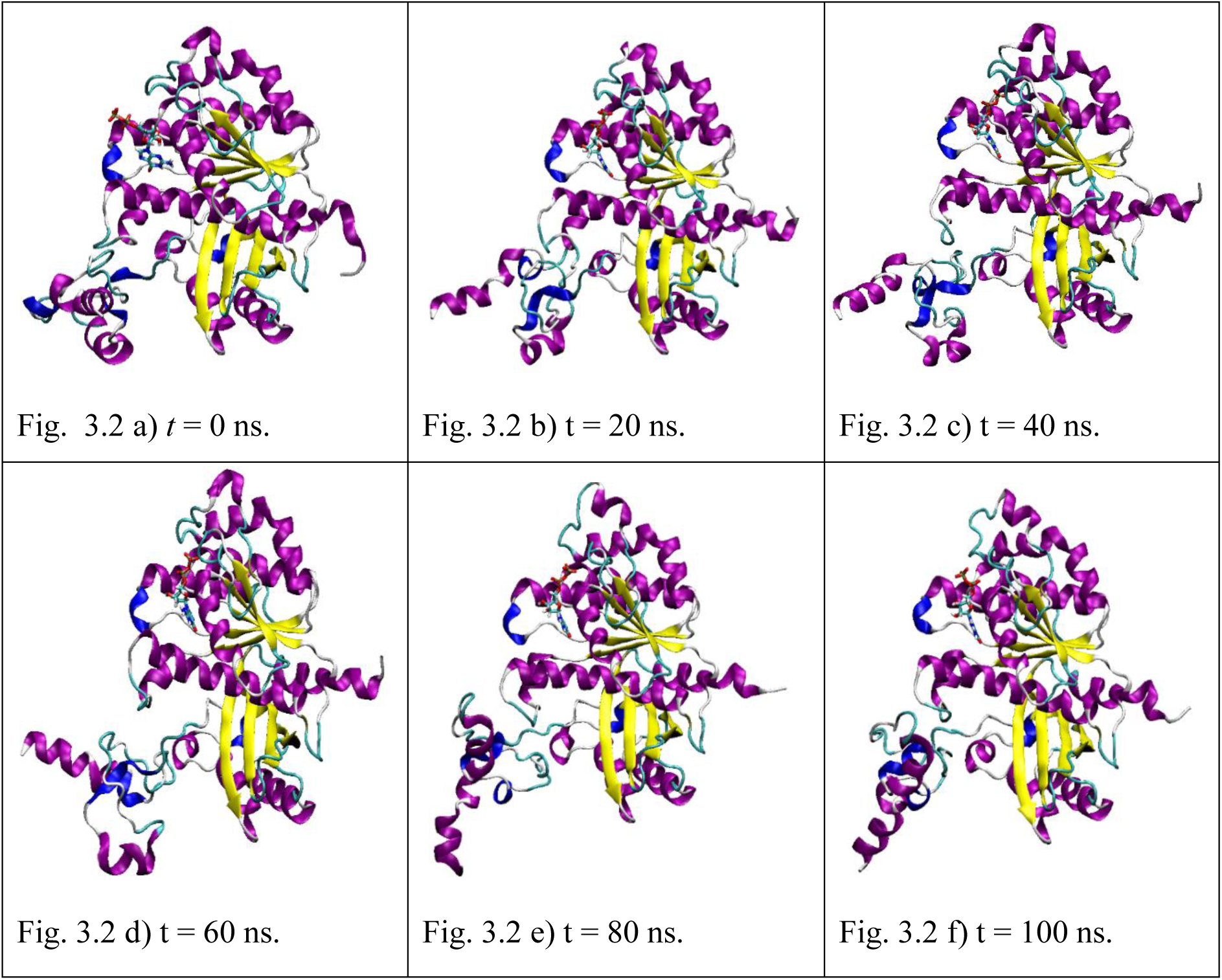
a – f) Structure of the protein in the wild-type monomer simulation WT-FtsZ mono-GDP (1) at every 20 ns from t = 0 ns to t = 100 ns during MD simulation: a) t = 0 ns, b) t = 20 ns c) t = 40 ns, d) t = 60 ns, e) t = 80 ns, f) t = 100 ns. The frames were aligned for the backbone atoms residues 12 – 311.

**Fig. 3.3.**
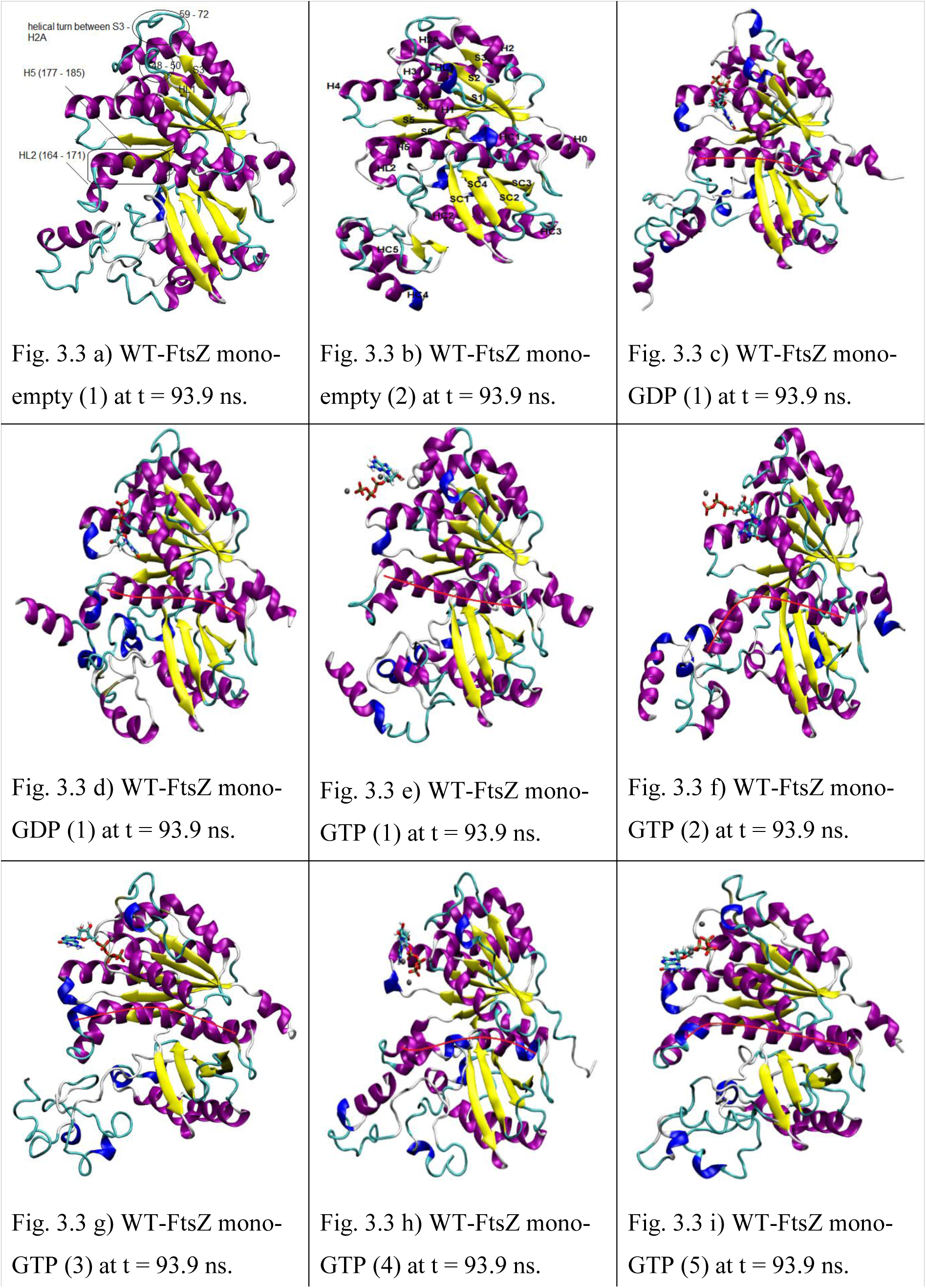
a – i) FtsZ structures at 93.9 ns simulation time. VMD ‘new cartoon’ representation is used for the protein and ‘secondary structure’ colour scheme is used. ‘CPK model’ representation is used for nucleotide and coloured based on atom name; and ‘CPK model’ for Mg^2+^ ion, coloured in black. To bending of the central helix is shown using a red line drawn through the helix.

The curvature of the central helix in the representative figures are shown using a red line along the helix. In the simulations with GTP, the residues 177 – 188 *i.e.* the N-terminal residues of the central helix bends towards the C-terminal domain, thus opening the nucleotide binding site considerably. This was observed in at least four out of the five simulations, WT-FtsZ mono-GTP (2), (3), (4) and (5) (Figure 3.3 (f) – (i)). In the simulation WT-FtsZ mono-GTP (1), the bend was observed for a short duration much before the 93.9 ns frame. In the simulations, WT-FtsZ mono-GTP (2) and WT-FtsZ mono-GTP (4) (Fig. 3.3 (f) and (h)) a break in the central helix is observed which divides it into two. This is similar to the kinked helix reported in the *S.aureus* FtsZ crystal structure apo form [29]. However, the overall structure of the globular core observed in the *S.aureus* is quite different in which the two β-planes appear to merge and form a single large plane.

On the other hand, in simulations with GDP (Fig. 3.3 d) the central helix may bend towards the N-terminal domain (Figure 3.3 d). There are no major distortions in the helix in GDP simulations (changes in secondary structure or break in helix).

It may be observed that residues of the HC2-SC2 loop (245 – 257), have varying conformations. In Fig. 3.3 (a), (b) only one 3-10 helix is present, in Fig. 3.3 (c), (d) and (e), (g) and (i) two 3-10 helices are observed, in Fig. 3.3 (f) and (h) an α-helix and a 3-10 helix are present.

### 3.4 Root mean square deviation (RMSD)

To study the stability of the model during molecular dynamics simulations, root mean square deviation (RMSD) of backbone atoms was calculated. RMSD measures the structural stability of the protein during the course of simulation and may be used to test if there are large structural changes in the protein leading to high RMSD. RMSD was calculated for the backbone atoms of residues 12 – 311 *i.e.* the globular core. The structure at time, t=0 *i.e* after npt-equilibration was used as reference for fitting. The trajectories were fit over backbone atoms of residues 12 – 311. RMSD was calculated at every 20 ps.

RMSD for simulations are provided in Figure 3.4 (a – c). A steady increase in RMSD during the initial 20 ns period was observed. This corresponds to the equilibration / stabilization phase of the simulation. After this equilibration phase, structural drifts in the core globular domain minimize, and a more or less stable RMSD is attained. RMSD fluctuates within ∼0.5 Å. This range is quite similar to RMSD reported in previous studies [6, 7] in which crystal structures were used in simulations.

**Fig 3.4.**
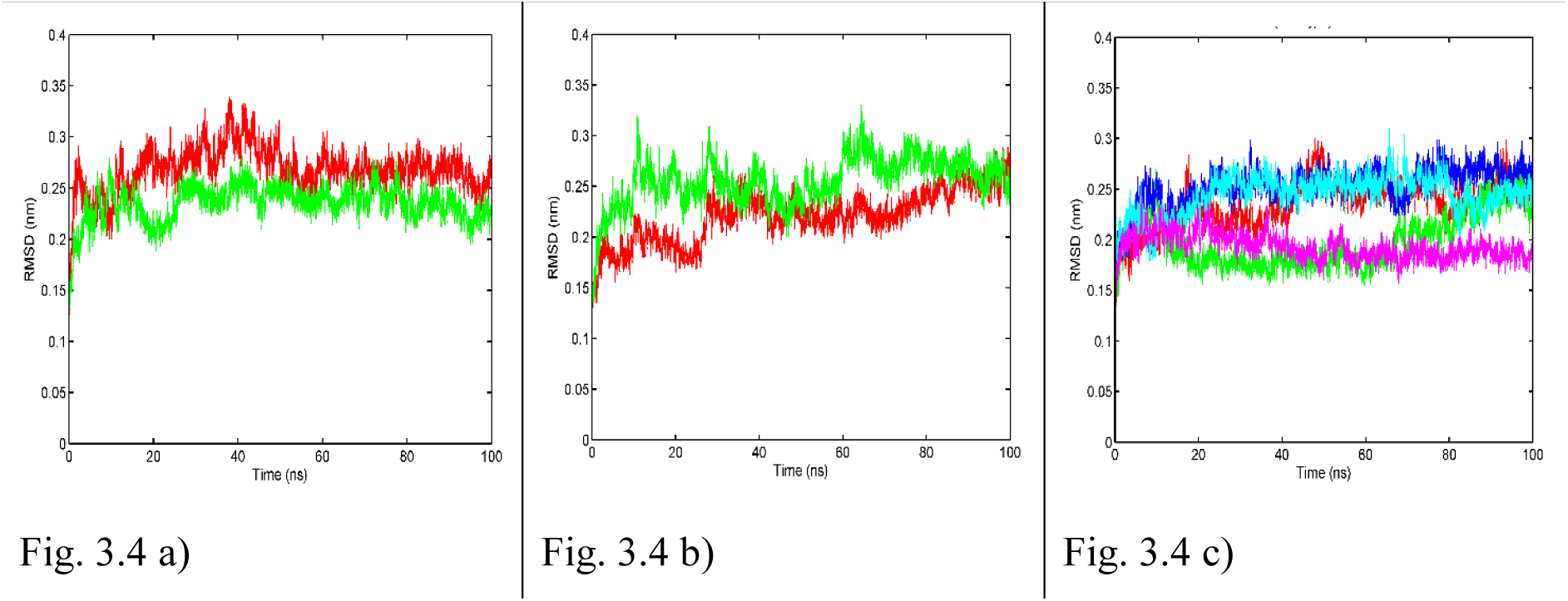
a – c) RMSD of backbone atoms of residues 12 – 311 of the globular tubulin like core of FtsZ in (a) WT-FtsZ mono-empty (1) (red) and WT-FtsZ mono-empty (2) (green). (b) WT-FtsZ mono-GDP (1) (red) and WT-FtsZ mono-GDP (2) (green). (c) WT-FtsZ mono-GTP (1) (red), WT-FtsZ mono-GTP (2) (green), WT-FtsZ mono-GTP (3) (blue), WT-FtsZ mono-GTP (4) (cyan), WT-FtsZ mono-GTP (5) (magenta). For RMSD calculation, the simulation trajectory frames were aligned on backbone atoms of residues 12 – 311 of the structure obtained after respective NPT equilibrations.

Important conclusions which may be drawn from the RMSD plot are 1) the globular core is stable and does not undergo any large structural changes and 2) the presence of CTL does not affect the structure or stability of the globular core.

### 3.5 Root mean square fluctuation (RMSF)

Average RMSF per residue was calculated for backbone atoms of residues 2 – 383 from 20 – 100 ns *i.e.* after equilibration (Fig 3.5). High RMSF of the N-terminal helix H0, is consistent with its intrinsic disorder predicted by PrDOS (Section 3.1). Residues of the loop HL1 have high flexibility. It may be observed that these residues 48 – 50 may take conformation of loop (Fig. 3.2 (a) and (f)), alpha helix (Figure 3.2 (d)) and 3-10 helix (Figure 3.2 (b), (c), (e), (h) – (i)). Residues of the loop between S3 and H2A, 59 – 72, may have high flexibility. The residues of helix HL2, loop HL2-H5 and few N-terminal residues of the central helix H5 have high flexibility (residues 165 – 175). This is consistent with the changes in secondary structure (α-helix to 3-10 helix) and bending observed for residues in this region of the protein. The residues in this region form contacts with HC3 of the upper subunit during polymerization [*M. jannaschii* FtsZ dimer 1W5A, 1W5B]. The bending of the central helix in simulations with GTP, makes the central helix more accessible for making protofilament contacts. Consistent with the intrinsic disorder of the IDR, RMSF for residue no. 320 onwards were much higher than the rest of the protein. Two to three major peaks are observed in this region 1) between 321 – 341, 2) near 361 and 3) the extreme C-terminal tail which has the maximum RMSF. In between the first two peaks of the IDR there is a sharp decline in RMSF. Such an RMSF profile indicates that there is some structural order due to the presence of less flexible residues within the IDR. Further, it may be observed that the maximum RMSF for residues in this region in simulations of FtsZ with GTP are less than 0.55 nm but in simulations with GDP it may go up to upto1.64 nm (0.8 nm in the other).

**Fig. 3.5).**
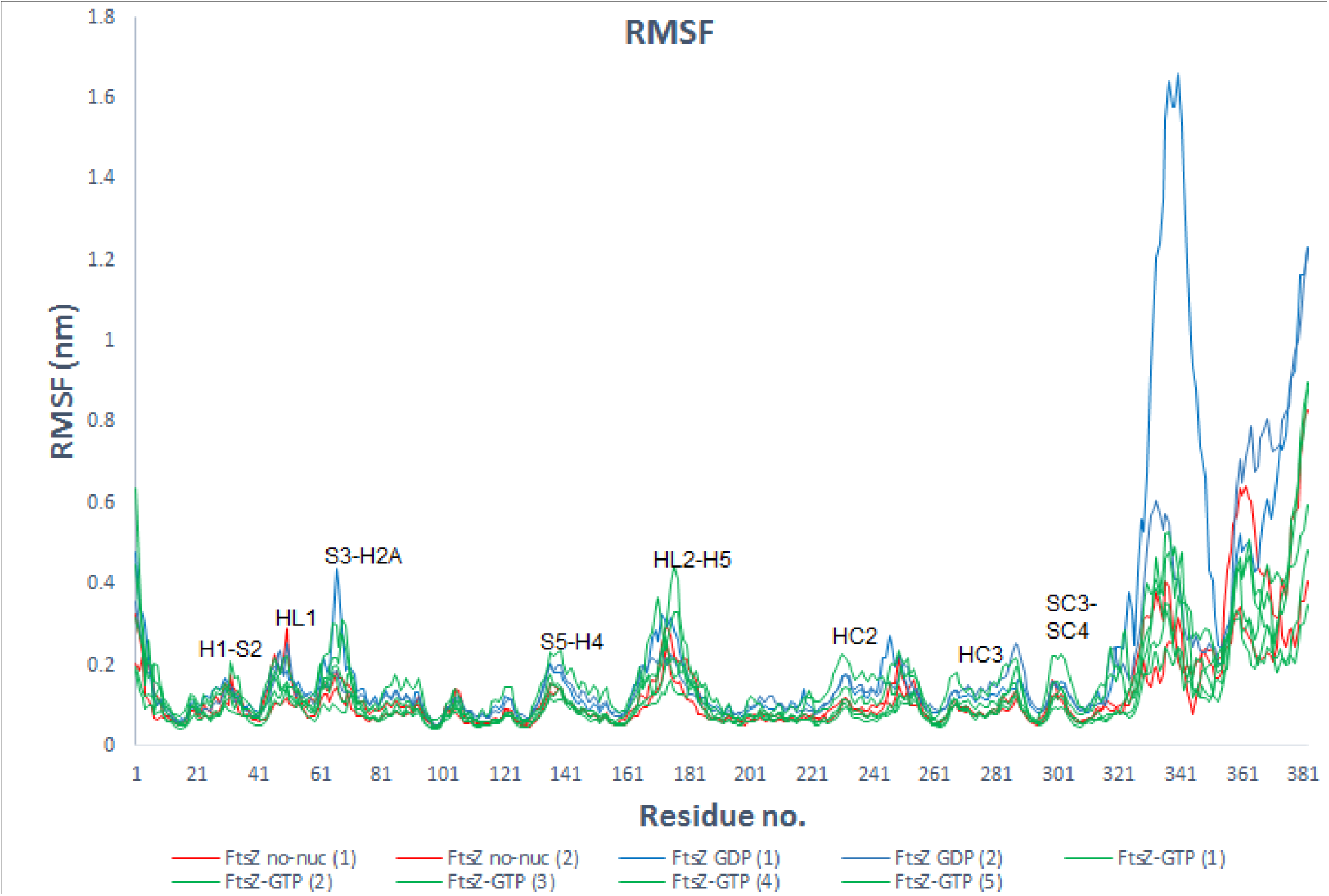
RMSF of protein backbone atoms and averaged per residue from 20 - 100 ns WT-FtsZ mono-empty (red); WT-FtsZ mono-GDP (blue) WT-FtsZ mono-GTP (green).

Since many interactions with regulatory proteins occurs at the CTL, a highly flexible CTL would increase interaction of the GDP bound monomer and regulatory proteins. In contrast a lower flexibility CTL, would decrease interaction with regulatory proteins, in turn allowing polymerization of the GTP form.

### 3.6 Hydrogen Bonds between nucleotide and protein

For calculating hydrogen bonds distance cut-off of 3.5 Å and angle cut-off 25° was used. Hydrogen bonds were calculated during 0 – 100 ns MD simulation. ‘Hydrogen bond occupancy’ is defined as the percentage time during which the hydrogen bond is present in the simulation. Hydrogen bonds which have occupancy greater than 38% are given in Table 3.2. In the simulation WT-FtsZ mono-GDP (1), GDP forms hydrogen bonds with Glu137, Arg141, Gly106 and Thr131. In the simulation WT-FtsZ mono-GDP (2), GDP forms 2 hydrogen bonds only with Glu137, Arg141. In the simulation WT-FtsZ mono-GTP (1), GTP forms hydrogen bonds with Arg141 and Lys140. In the simulation WT-FtsZ mono-GTP (2), GTP forms hydrogen bonds with Arg141, Lys140 and Glu137. In the other three simulations, GTP forms hydrogen bonds with Arg141 only.

**Table 3.1.** Hydrogen bonds between nucleotide and protein.

| S.No. | Simulation | Hydrogen Bond with nucleotide | Occupancy (%) |
| --- | --- | --- | --- |
| 1 | WT-FtsZ mono-GDP (1) | GLU137-Side | 95 |
|  |  | ARG141-Side | 71 |
|  |  | GLY106-Main | 48 |
|  |  | THR131-Side | 45 |
| 2 | WT-FtsZ mono-GDP (2) | GLU137-Side | 84 |
|  |  | ARG141-Side | 66 |
| 3 | WT-FtsZ mono-GTP (1) | ARG141-Side | 75 |
|  |  | LYS140-Side | 68 |
| 4 | WT-FtsZ mono-GTP (2) | ARG141-Side | 63 |
|  |  | LYS140-Side | 61 |
|  |  | GLU137-Side | 61 |
| 5 | WT-FtsZ mono-GTP (3) | ARG141-Side | 86 |
| 6 | WT-FtsZ mono-GTP (4) | ARG141-Side | 82 |
| 7 | WT-FtsZ mono-GTP (5) | ARG141-Side | 68 |

**Table 3.2.** Contacts between the IDR and the globular core (Contact maps provided in SI)

| Simulation | Contacts with HL2-H5 | Contacts with C-terminal domain |
| --- | --- | --- |
| WT-FtsZ mono-empty (1) | 351 – 355 and 168 – 171 | 339 – 342 and 228 –233 |
| WT-FtsZ mono-empty (2) |  | 342 – 346 and 245 –249 |
| WT-FtsZ mono-GDP (1) |  | 353 – 358 and 246 – 252 |
| WT-FtsZ mono-GDP (2) | 349 – 354 and 169 – 172 | 355 – 357 and 246 – 248 |
| WT-FtsZ mono-GTP (1) | 347 – 350 and 169 – 171 | 354 – 356 and 245 – 248 |
| WT-FtsZ mono-GTP (2) | 350 – 355 and 167 – 172 | 355 – 358 and 247 – 249 |
| WT-FtsZ mono-GTP (3) | 352 – 354 and 168 – 173 | 347 and 238 – 240 |
| WT-FtsZ mono-GTP (4) | 348 – 352 and 168 – 173 |  |
| WT-FtsZ mono-GTP (5) | 352 – 355 and 168 – 171 | 353 – 356 and 247 – 253 |

The residues Arg141, Lys140 and Glu137 belong to the S5-H4 loop. Since both GTP and GDP bind to the same S5-H4 loop residues and GTP can bind from various positions from the outside by forming non-specific hydrogen bonds (Fig. 3.3 (e) and (h)), i*n vivo*, introduction of GTP in the nucleotide binding site of the FtsZ-GDP monomer would enable release of GDP. The bending of the central helix would also enable the release of GDP.

### 3.7 Principal Component Analysis

Principal component analysis was performed to identify significant concerted motions in different regions of the protein and understand protein dynamics. Covariance analysis was performed for the Cα atoms of the monomer protein. Filtered trajectories for the Cα atoms were generated for the first eigen vector (Fig. 3.6).

**Figure 3.6.**
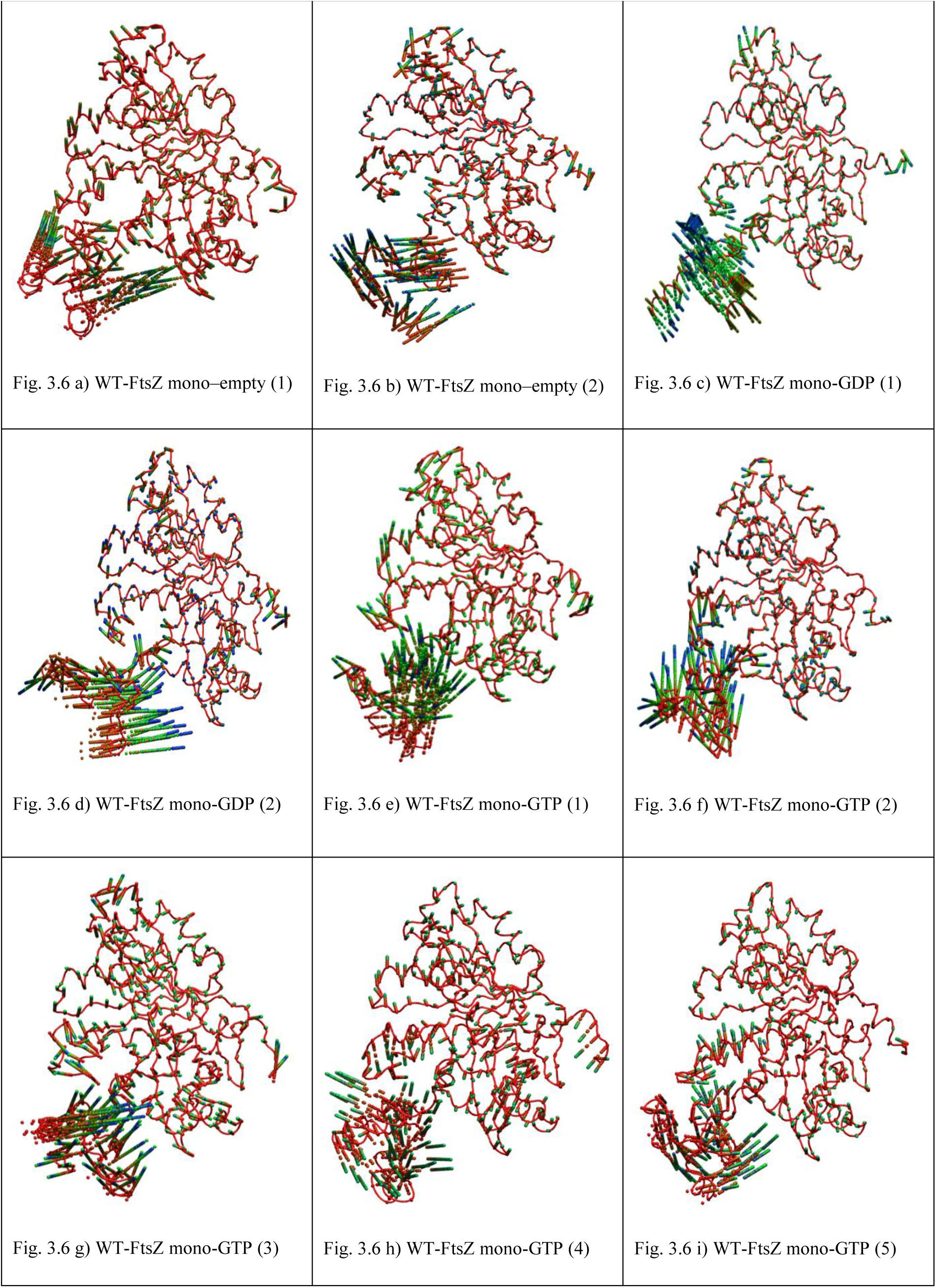
a – i) Principal motion along the first eigen vector for (a – b) WT-FtsZ-empty (1) and (2), (c – d) WT-FtsZ-GDP (1) and (2) and (e – i) WT-FtsZ-GTP (1), (2), (3), (4) and (5). The position of atoms at different simulation timesteps is shown using the ‘CPK’ representation in VMD. The ‘trajectory-timestep’ colour scheme in VMD has been used with the ‘RGB’ colour scale such that the first frame (*t* = 100 ps MD simulation time) is in red and the frame at *t* = 100,000 ps is in blue. In addition, the first frame is also shown using the ‘Tube’ representation in VMD so that the protein / the secondary structure elements may be visualized.

It may be observed that the Cα atoms of the helix H0, HL1, the S3 – H2A loop, the S5 – H4 loop, helix HL2, the HL2 – H5 loop, the N-terminal residues of the central helix H5, the loop SC1 – HC2, helix HC2, loop HC2 – SC2, HC3 and the loop SC3 – SC4 have higher displacements among the Cα atoms of the residues 1 to 312 *i.e.* the N-terminal helix H0 and the ordered residues of the N- and C-terminal.

The higher displacements of H0 is expected due to its intrinsic disorder. The nucleotide binding site residues HL1, the S3 – H2A loop, the S5 – H4 loop, helix HL2, the HL2 – H5 loop, the N-terminal residues of the central helix H5 have high displacements. The secondary structure changes were described in Section 3.3 and 3.5.

Rotation of the C-terminal domain may be observed in Fig. 3.6 (a) and (b) (downwards) and (e), (g) and (h) (upwards). In GDP bound simulations, the magnitude is less (compared to the two empty- and three GTP bound simulations Fig. 3. 6 (e), (g) and (h)). Between the three GTP bound simulations, in which rotation is observed, the domain motion is towards the front in (e), towards the nucleotide binding site in (g) and more perpendicular in (h). The displacement of the Cα atoms of the C-terminal flexible or disordered region are much higher than the rest of the protein.

Average Cα coordinates for the ordered residues were compared after alignment of N-terminal ordered residues (12 – 164) (Fig. 3.7). It was observed that the C-terminal domain is rotated or shifted more towards the left (towards the nucleotide binding site) in simulations with GTP. In contrast in the simulations with GDP, the domain is shifted towards the right.

**Fig. 3.7).**
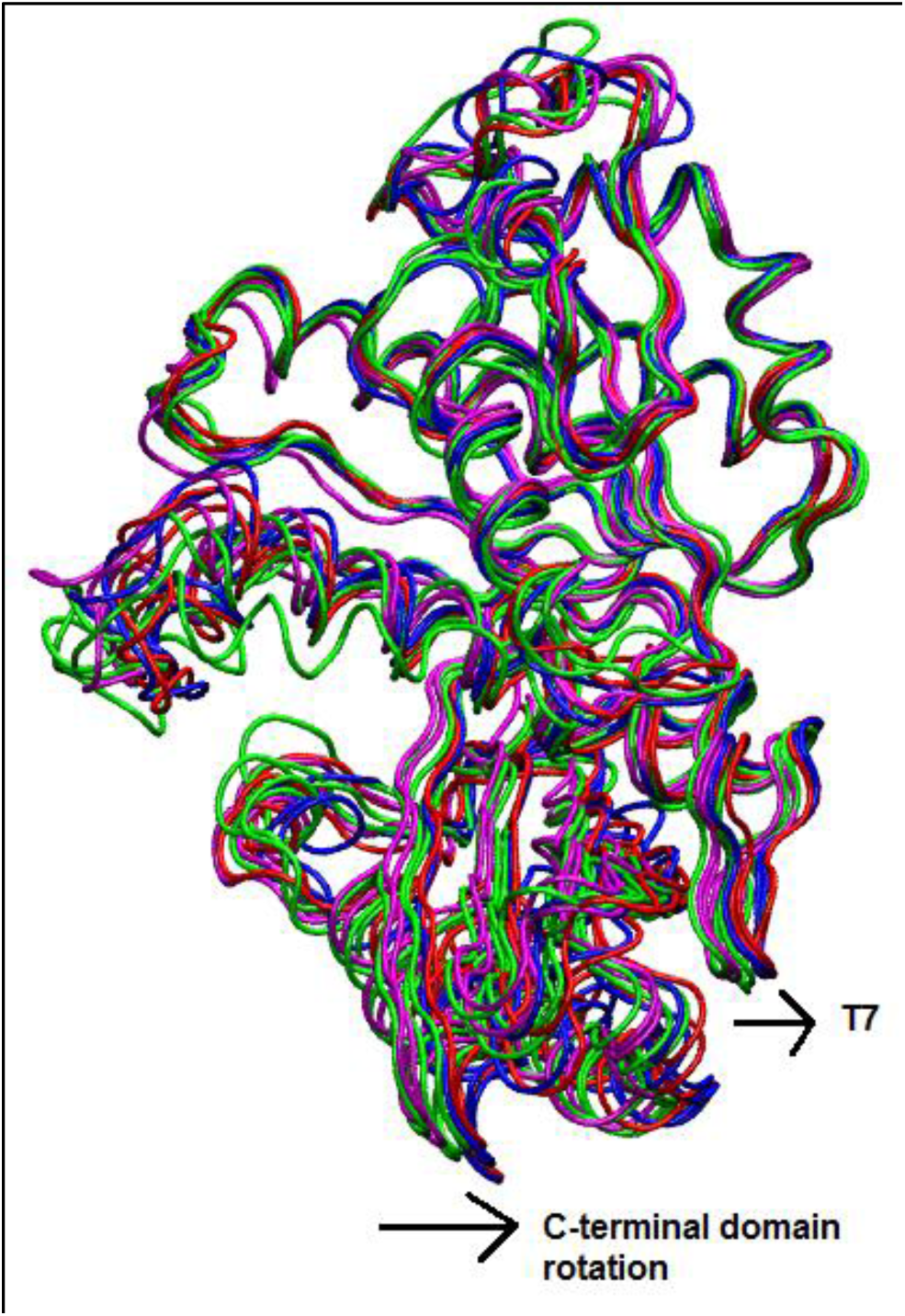
Alignment of average Cα coordinates after superpositioning Cα atoms of N-terminal residues 12 – 164: WT-FtsZ mono-empty simulations (blue), WT-FtsZ mono-GDP simulations (red), WT-FtsZ GTP simulations (magenta and green).

## 4. Conclusions

FtsZ monomer was simulated without nucleotide, with GTP and with GDP. The *E. coli* FtsZ protein was simulated. The structure for the protein was constructed by homology modelling. It also includes the C-terminal IDR. The protein model was found to be stable during simulations. The structural stability of the ordered domains was found to be comparable to what was reported in previous studies. The IDR has high structural variability. Residues with much lower flexibility are present within the IDR. Such residues with low RMSF provide some 3-D form to the IDR (it is not a loose open chain of residues). In the simulations with GDP, the IDR is more flexible than in simulations with GTP or without nucleotide. It was observed that GTP binds from the outside of the nucleotide binding site, non-specifically to the long chain residues of the S5-H4 loop (Arg141, Lys140, Glu137). A significant opening of the nucleotide binding site was observed in simulations with GTP which is caused by the bending of the core helix. Such a bending of the central helix, would facilitate nucleation polymerization to occur. The structure of nucleus being the GTP bound FtsZ monomer to which other GTP bound monomers may add easily. It was proposed that the CTL has an important role in transforming FtsZ from an inactive to an active state [8, 9] by providing structural flexibility to the N-terminal core and in subunit subunit interaction.

Principal component analysis suggested concerted motion of the Cα atoms of helix H0, the S3 H2A loop, the S5 – H4 loop, the helix HL2, the HL2 – H5 loop, the N-terminal residues of the central helix H5, the loop SC1 – HC2, the helix HC2, the helix HC3, the loop SC3 – SC4 and the IDR. In the simulations of the *A. aeolicus* FtsZ monomer [6], concerted motion was observed for residues in the HL2 – H5 loop, the S3 – H2A loop, the H5 – HC1 loop, the helix HC2 and HC3. However, it was observed that in the GTP/GDP bound simulations, HC2 and HC3 move downwards (diagonally opposite from the nucleotide binding site in the GTP bound state *i.e.* towards the adjacent monomer of the protofilament). In our simulations, we observed that in all the simulations with GTP, the domain shifts forward, upwards or diagonally towards the nucleotide binding site (Fig. 3.6 (e), (g) and (h)). Therefore, our observation suggests that the GTP bound FtsZ favours polymerization by the bending of the central helix, allowing contacts to be formed at the N-terminal domain but the C-terminal domain shifts upwards in the monomer.

The differential flexibilities observed in the presence of GTP and GDP supports a regulatory role for the IDR. In the absence of nucleotide or presence of GTP higher up in the nucleotide binding site, results in a relatively lower flexible IDR. In contrast, in simulations with GDP, the flexibility is much higher. Our simulations suggest that variable/non-specific GTP binding to the monomer is a key feature of FtsZ dynamics. In the presence of a well bound stable nucleotide, the central helix may be stabilized by interactions with nucleotide but in the presence of GTP bound non-specifically, the monomer first allows polymerization to occur at the nucleotide binding site. Contact maps from the average Cα coordinates were generated to find contacts between residues of the IDR (316 to 382) and residues 164 – 310, HL2 to SC4. Usually 2 – 3 contacts are formed between residues in 341 – 361 and 1) in HL2-H5 2) in HC2, HC2-SC2 loop. For e.g. in the simulation WT-FtsZ mono-GTP (1), 2 contacts are observed, between 347 – 350 and 169 – 171 (in HL2-H5), and between 354 – 356 and 245 – 248 (in HC2-SC2 loop) (Table 3.2). Consistent with this, a low RMSF region is present between residues 341 – 361 in all simulations. By performing sequence analysis in VMD, it was observed that in residues 341 – 361, there are 10 non polar residues (47 %); in 164 – 177, there are 6 non-polar residues (43%) and in 243 – 253, there are 6 non-polar residues (54%). Since this region forms contacts with the HL2-H5 loop and the ordered C-terminal domain, it is possible that the C-terminal IDR may be directly involved in bending of the core helix and rotation of the C-terminal domain through hydrophobic interactions. In Fig. 3.7, average structures show a large difference in residues of the HC2–SC2 region (243 – 253). PCA analysis also shows significant displacements of these residues (Fig. 3.6 (a), (c), (f), (i)). A twisted bending of the central helix in the nucleotide free simulation of *A.aeolicus* FtsZ monomer was observed in the simulation without nucleotide [6]. Super positioning of average Cα coordinates of their simulations showed that in the same simulation the C-terminal domain shifts towards the nucleotide binding site (similar to the GTP bound simulations of the *E.coli* FtsZ). Therefore, it appears that hydrophobic interaction between residues of HL2–H5 (164 – 177) and the helix HC2, HC2– SC2 exists which causes rotation in the nucleotide free simulation of *A.aeolicus* FtsZ [6]. However, in the presence of GTP/GDP they do not observe such bending of the central helix, similar to our GDP bound simulations of *E.coli* FtsZ.

In our simulations, in the nucleotide free form, the hydrophobic interaction between HL2-H5 and HC2-SC2 is limited by steric interactions with the IDR. In the presence of GDP, the central helix is stabilized by the nucleotide. Due to its higher charge, GTP is not able to bind stably in the nucleotide binding site. Since it forms hydrogen bonds from the outside or higher up in the nucleotide binding site with the long chain charged residues of the S5-H4 loop (Arg141, Lys140, Glu137), it may lead to differential charge distribution in the nucleotide binding site which allows the IDR to pull the central helix downwards even in the presence of GTP.

Our simulations support recent experiments in which the presence of IDR was found to be essential for FtsZ polymerization [8, 9]. In the experiments it was found that in the absence of the C-terminal linker, FtsZ critical concentration increases, however GTPase activity reduces from 3.82 GTP FtsZ^−1^min^−1^ to 1.08 GTP FtsZ^−1^min^−1^, protofilaments were observed after a long delay (15 min after addition of GTP) [8]. From our simulations we may say that in the absence of the IDR, few nucleus FtsZ would be available to allow polymerization. When the IDR residues are scrambled, then the distribution of non-polar residues is affected, due to which a lower GTPase activity compared to the wild-type is seen in the experiments.

In conclusion, we may say that the IDR is not an open chain of amino acids, it has some structural order within to allow interaction with the globular core. We observed that the C-terminal IDR forms hydrophobic contacts with the HL2-H5 residues and the HC2, HC2-SC2. Through such interactions with the globular core, it is able to bend the core helix in the GTP bound FtsZ monomer, favouring polymerization.

## Acknowledgements

We would like to thank the Marie Curie Actions program for funding the project, supercomputing resources at the Warwick University and the University of Southampton, Dr. Alison Rodger and Dr. Syma Khalid for their supervision in the project.

